# Replicative stress in gastroesophageal adenocarcinoma is associated with chromosomal instability and sensitivity to DNA damage response inhibitors

**DOI:** 10.1101/2023.03.27.534412

**Authors:** Pranshu Sahgal, Deepa T. Patil, Zsofia M. Sztupinszki, Viktoria Tisza, Sandor Spisak, Brandon Huffman, Aurel Prosz, Harshabad Singh, Jean-Bernard Lazaro, Zoltan Szallasi, James M. Cleary, Nilay S. Sethi

**Affiliations:** Department of Medical Oncology, Dana-Farber Cancer Institute, Boston, MA, 02215, USA; Department of Medicine, Brigham and Women’s Hospital and Harvard Medical School, Boston, MA, 02115, USA; Broad Institute of Massachusetts Institute of Technology (MIT) and Harvard University, Cambridge, MA, 02142, USA; Department of Pathology, Brigham and Women’s Hospital, Harvard Medical School, Boston, MA 02215, USA; Danish Cancer Society Research Center, Copenhagen, 2100, Denmark; Department of Radiation Oncology, Dana-Farber Cancer Institute, Boston, MA, 02215, USA; Center for DNA Damage and Repair (CDDR), Dana-Farber Cancer Institute, Boston, MA, 02215, USA; Computational Health Informatics Program, Boston Children’s Hospital, Boston, MA, 02115, USA; Division of Gastrointestinal Oncology, Department of Medical Oncology, Dana-Farber Cancer Institute, Boston, MA, 02215, USA

**Keywords:** Gastric cancer, esophageal cancer, DNA-damage response, aneuploidy, replication stress, CIN, DNA damage response pathway inhibitors, irinotecan

## Abstract

Gastroesophageal adenocarcinoma (GEA) is an aggressive, often lethal, malignancy that displays marked chromosomal instability (CIN). To understand adaptive responses that enable CIN, we analyzed paired normal, premalignant, and malignant gastric lesions from human specimens and a carcinogen-induced mouse model, observing activation of replication stress, DNA damage response (DDR), and cell cycle regulator p21 in neoplastic progression. In GEA cell lines, expression of DDR markers correlated with ploidy abnormalities, including high-level focal amplifications and whole-genome duplication (WGD). Moreover, high expression of DNA damage marker *H2AX* correlated with CIN, WGD, and inferior patient survival. By developing and implementing a composite diagnostic score that incorporates *TP53* mutation status, ploidy abnormalities, and *H2AX* expression, among other genomic information, we can identify GEA cell lines with enhanced sensitivity to DDR pathway inhibitors targeting Chk1/2 and Wee1. Anti-tumor properties were further augmented in combination with irinotecan (SN38) but not gemcitabine chemotherapy. These results implicate specific DDR biomarkers and ploidy abnormalities as diagnostic proxy that may predict premalignant progression and response to DDR pathway inhibitors.

## INTRODUCTION

Gastroesophageal adenocarcinoma (GEA) is a leading cause of death globally and improvement in patient outcomes has been hampered by multiple issues, including failure to identify patients with high-risk precancerous lesions and limited targeted therapy options for patients with advanced disease. The Cancer Genome Atlas project’s (TCGA) recent molecular classification has characterized gastroesophageal adenocarcinoma into 4 subtypes: microsatellite-unstable, Epstein-Barr virus-positive, chromosomal-instability (CIN), and genomically stable cancers. These classifications have given insight into the molecular features of these subtypes, helping advance the field of early diagnosis and targeted therapies in GEA(Cancer Genome Atlas Research, 2014).

The most common GEA subtype is CIN, seen in 50% of patients. The mechanisms leading to and consequences of CIN are still poorly understood(Maleki and Rocken, 2017; Sohn et al., 2017). Nevertheless, CIN is positively correlated with multidrug resistance and ploidy abnormalities, including whole-genome duplication (WGD) and high-level focal amplification events. In addition, recent studies have hypothesized that replicative stress, defined as the presence of stalled replication forks leading to single-strand DNA and possible fork collapse, plays a major role in CIN development. Several proteins in the DNA damage response (DDR) pathway, including Chk1 and Wee1, compensate for replicative stress by slowing the cell cycle and promoting the stability of stalled replication forks(Dominguez-Kelly et al., 2011; Elvers et al., 2012). When these compensatory mechanisms fail, replication forks can collapse and lead to the generation of double-strand DNA breaks, promoting CIN.

We hypothesized that high levels of replicative stress are present in GEA CIN oncogenesis. We evaluated the expression of DDR markers in human and mouse gastric precancer and malignant specimens. Further, we assessed whether the GEA cell lines with different levels of DDR and associated ploidy abnormalities correlate with sensitivity to DDR inhibitors in gastric cancer. Recent preclinical and clinical studies suggest that the chemotherapeutic agents that generate replication stress synergize with Chk1 inhibitors across cancers(Banerji et al., 2019). Therefore, we also examined anti-tumor properties of DDR inhibitors in combination with two commonly used chemotherapeutic agents, showing greater therapeutic potential for one pair.

## RESULTS

### Activation of select DDR markers in human gastric premalignancy progression

The progression from premalignancy to cancer is associated with replication stress that leads to DNA double-strand breaks (DSBs) and genomic instability(Macheret and Halazonetis, 2015). To characterize DDR/replication stress in GEA premalignancy progression, we procured human gastric specimens from 8 patients that had matched normal, premalignant, and malignant lesions (Table S1). Using immunohistochemistry (IHC), we evaluated the levels of H2AX/phospho-H2AX (pH2AX) and 53BP1, 2 well-known markers of DNA DSBs and replicative stress(Bartkova et al., 2005; Gupta et al., 2014; Mah et al., 2010). We observed that 53BP1 levels increased during neoplastic progression, with invasive adenocarcinoma displaying the highest levels for nearly all patients (Figure 1A-C and Table S2). H2AX and pH2AX were largely absent from normal stomach and premalignant lesions, but malignant lesions from 2 patients showed moderate staining of H2AX and pH2AX (Figure 1A-B, Figure S1A and Table S2). These observations suggest that human gastric premalignant to malignant progression is associated with DNA DSBs and replication stress.

**Figure 1:**
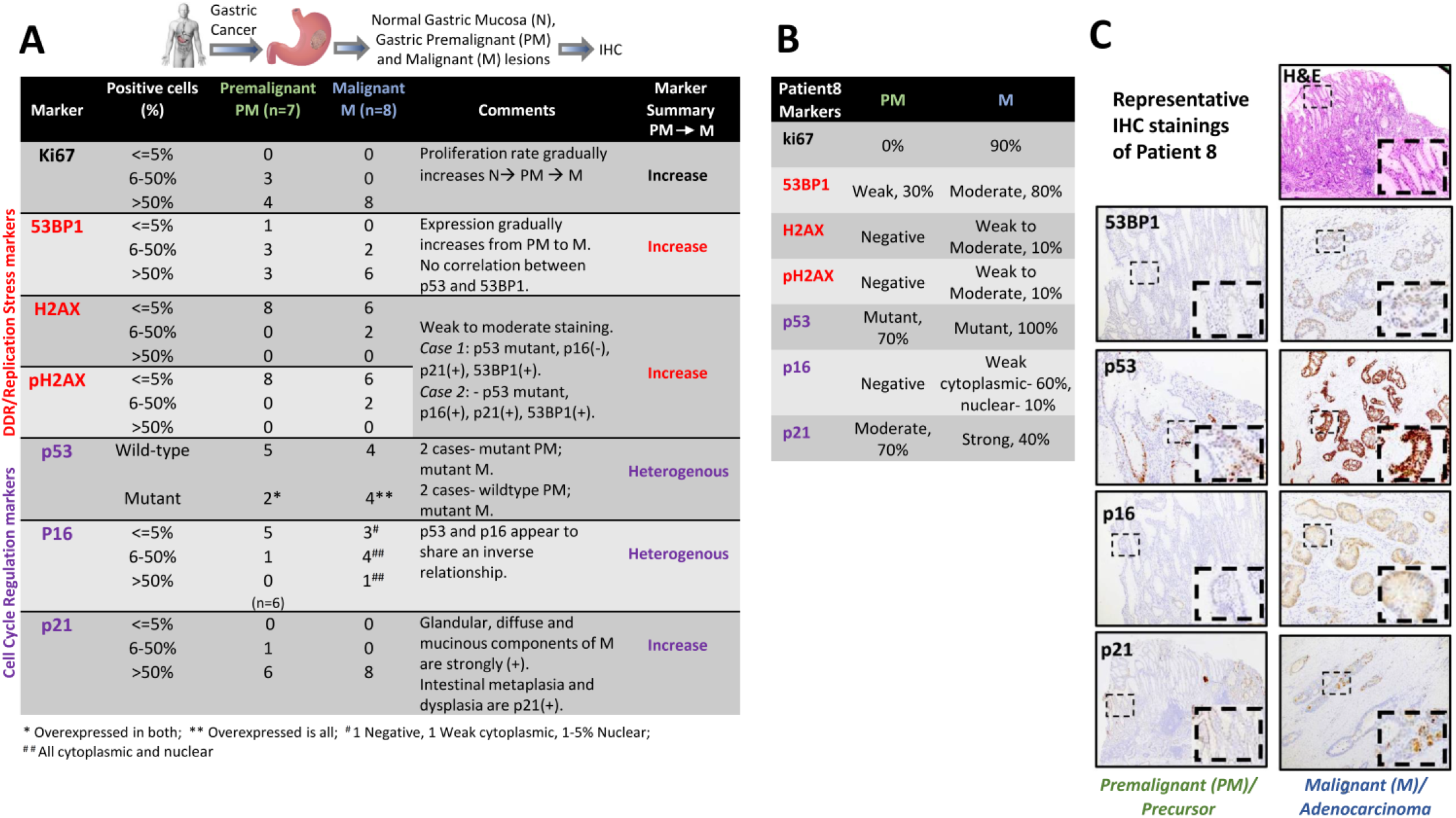
Evaluation of DDR and cell cycle markers in human gastric specimens. A. Tabulated summary of IHC stainings indicating the number of premalignant (green) and malignant (blue) gastric lesions scored for each staining criterion. DDR/replication stress markers-53BP1, H2AX, pH2AX in red; Cell cycle regulators-p53, p16, p21 in purple. B. Summary of the stainings (DDR markers and the cell cycle regulation markers) in paired premalignant and malignant gastric lesions from gastric cancer patient8. PM= Premalignant gastric lesion, M= Malignant gastric lesion. C. Representative IHC images of paired premalignant and malignant gastric lesions from a gastric cancer patient (patient8) stained for DDR markers and cell cycle regulation markers, including H&E staining. Insets represent the zoomed area for each image.

Unlike colorectal cancer, GEA harbor early *TP53* mutations, often preceding dysplasia (Sethi et al., 2020; Stachler and Bass, 2020; Stachler et al., 2018; Stachler et al., 2015; Weaver et al., 2014). As GEA evolves, CIN ensues, selecting for amplification of oncogenes and deep deletions of tumor suppressor genes(Liu et al., 2018; Sahgal et al., 2021). We recently showed that *TP53*(p53)/*CDKN2A*(p16) co-altered premalignant gastric organoids derived from a mouse model displayed evidence of early CIN, which was associated with the activation of replication stress and DDR pathways(Sethi et al., 2020). Based on this understanding, we next examined the status of cell cycle regulation markers (p53, p16, and p21) in our samples. We observed marked heterogeneity in p53 and p16 levels among premalignant and malignant lesions, even within a patient. Lesions from the same patient expressed both wild-type (WT) and mutant p53 as well as areas of p16 positivity and negativity (Figure 1A). Interestingly, mutant p53 and p16 expression appeared to share an inverse relationship, especially in malignant lesions; p53 WT cells displayed p16 expression whereas p53 mutant cells were p16 negative (Figure 1A-C, Figure S1B, and Table S3). A downstream effector of p53, p21 induces cell cycle arrest (Chen, 2016). Notably, we observed a gradual increase in p21 levels in GEA progression; malignant lesions demonstrated significantly higher expression of p21 compared to normal and premalignant lesions in almost all the patients. The glandular, diffuse, and mucinous components of adenocarcinoma showed strong p21 staining, while intestinal metaplasia and dysplasia showed moderate levels (Figure 1A-C, Figure S1B, and Table S3). p21 staining appeared to be independent of mutant p53 expression in gastric cancer progression (Figure 1B-C, Figure S2B, and Table S3). These results indicate that p53 and p16 expression is heterogenous within lesions and through cancer progression, whereas p21 gradually increases from normal to malignant lesions independent of p53 status.

### Evaluation of DDR markers in carcinogen-induced mouse model of gastric premalignancy

We next evaluated our panel of biomarkers in a carcinogen-induced (MNU) mouse model of gastric premalignancy and cancer to validate our human observations. Mice exposed to drinking water containing MNU developed premalignant and malignant gastric lesions after 1 to 1.5 years(Sethi et al., 2020). We examined 4 mice treated with MNU that developed varying degrees of dysplasia and cancer (Figure 2A-B), staining formalin-fixed tissue for markers of dsDNA breaks, DDR pathway, and cell cycle regulators. Technical immunostaining limitations precluded interpretation of slides stained for H2AX (dsDNA breaks), 53BP1 (DDR pathway), and p16 (cell cycle). As expected, premalignant and malignant lesions demonstrated greater proliferation compared to normal gastric tissue by Ki67 IHC. We also found the levels of pH2AX, p53, and p16 expression were similar in our mouse lesions compared to the human specimen (Figure 2A-B and S2A); pH2AX was expressed in a premalignant gastric lesion from one mouse and a malignant lesion from another moue (mouse 3 and 4), but not in normal gastric tissue from any mice. While p53 expression was low in normal gastric tissue, premalignant lesions in mouse 3 and 4 displayed higher p53 expression, consistent with it being mutated. Similarly, p21 was not expressed in normal gastric tissue but was expressed to varying degrees in premalignant and malignant lesions in all 4 mice, with the strongest heterogeneous staining in mouse 3 and 4 (Figure 2A-B and S2A). These results indicate that, like human gastric premalignancy progression, a carcinogen-induced mouse model of gastric adenocarcinoma progression displays increased levels of DDR marker pH2AX and cell cycle regulator p21, with p21 expression independent of p53 mutation status.

**Figure 2:**
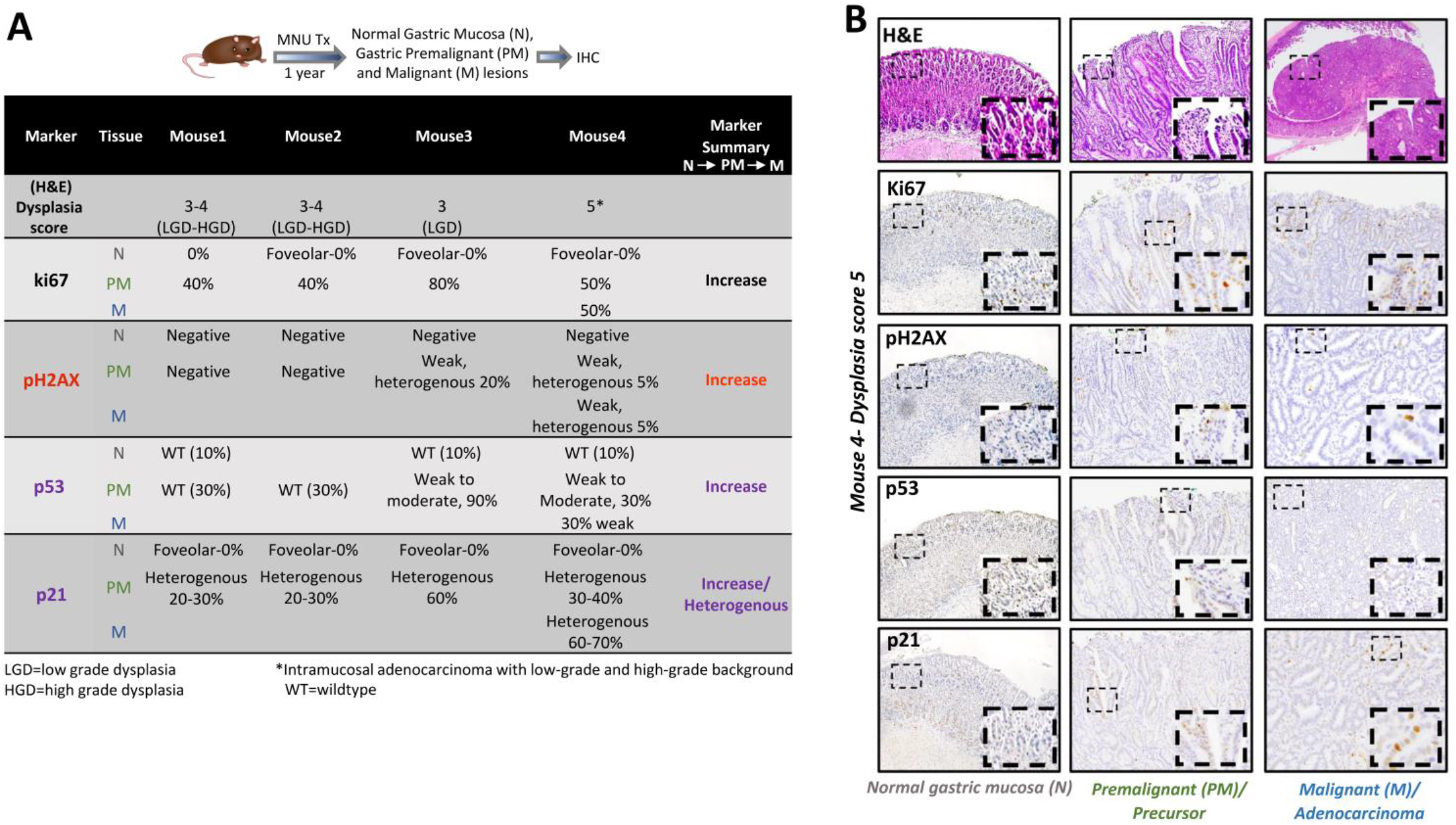
Evaluation of DDR and cell cycle markers in a carcinogen-induced mouse model of gastric premalignancy and cancer. A. Summary of dysplasia grade and staining pattern of proliferation marker ki67, DDR marker pH2AX, and cell cycle regulation markers (p53 and p21) in paired normal (grey), premalignant (green), and malignant (blue) gastric lesions of four MNU treated mice. B. Representative IHC images of the staining pattern of proliferation marker ki67, DDR marker pH2AX, and cell cycle regulation markers (p53 and p21) in paired normal, premalignant, and malignant gastric lesions of MNU treated mouse4 (dysplasia score 5).

### Evaluation of DDR markers in TCGA cohort of stomach adenocarcinoma

To further characterize components of the DDR pathway in human gastric cancer, we evaluated *H2AX* and *TP53BP1* expression in the TCGA stomach adenocarcinoma (STAD) dataset. We observed higher expression of both *H2AX* and *TP53BP1* in gastric adenocarcinoma samples compared with normal gastric tissue (Figures S2B). Interestingly, *H2AX* but not *TP53BP1* expression was significantly higher in *TP53* mutant gastric adenocarcinomas compared to *TP53* WT tumors (Figure S2C). Next, we evaluated the expression of *H2AX* and *TP53BP1* in Lauren’s classification subtypes of gastric adenocarcinoma. Lauren’s criteria is based on distinct clinical and molecular characteristics that divide gastric adenocarcinoma into intestinal and diffuse subtypes and is the most widely-used histological classification for gastric adenocarcinoma(Lauren, 1965). We observed a significantly higher expression of *H2AX* and not *TP53BP1* in the intestinal subtype than in the diffuse subtype (Figure S2D). Furthermore, *TP53* mutations were more enriched in intestinal subtype patients and in CIN tumors (Figure S2E). These observations suggest the patients with gastric adenocarcinoma that harbor p53 mutations and/or intestinal subtype histology express higher levels of DDR markers such as *H2AX*.

### DDR pathway activity is associated with ploidy abnormalities in gastric cancer

Since the majority of GEA exhibit ploidy defects as a result of CIN(Maleki and Rocken, 2017), we next evaluated the relationship between CIN and elevated markers of DDR or replication stress. To assess CIN in gastric cancer cell lines, we utilized the CIN25 score(Carter et al., 2006), a gene expression-based validated signature of CIN that predicts clinical outcomes across different human cancers. Indeed, gastric cancer cell lines in the Cancer Cell Line Encyclopedia (CCLE)(Barretina et al., 2012) displayed a strong correlation between the CIN25 score and a recently reported computational metric of aneuploidy (score based on Cohen-Shahir et al., 2021(Cohen-Sharir et al., 2021)) (Figure S3A). Importantly, *H2AX* expression correlated with high CIN25 score in gastric cancer cell lines (Figure 3A). In our analysis, we also found a positive correlation between the aneuploidy score and high-level focal amplifications (Figure S3B), a unique characteristic of gastric cancer evolution(Sahgal et al., 2021), a ploidy score (Figure S3C) and whole genome doubling (WGD) (Figure S3D) (both scores based on Ghandi et al., 2019(Ghandi et al., 2019)). Interestingly, gastric cancer cell lines with WGD have significantly higher expression of *H2AX* compared to ones without WGD (Figure 3B). These analyses suggest that DDR pathway markers, at least *H2AX*, track with WGD and ploidy defects in gatric cancer.

**Figure 3:**
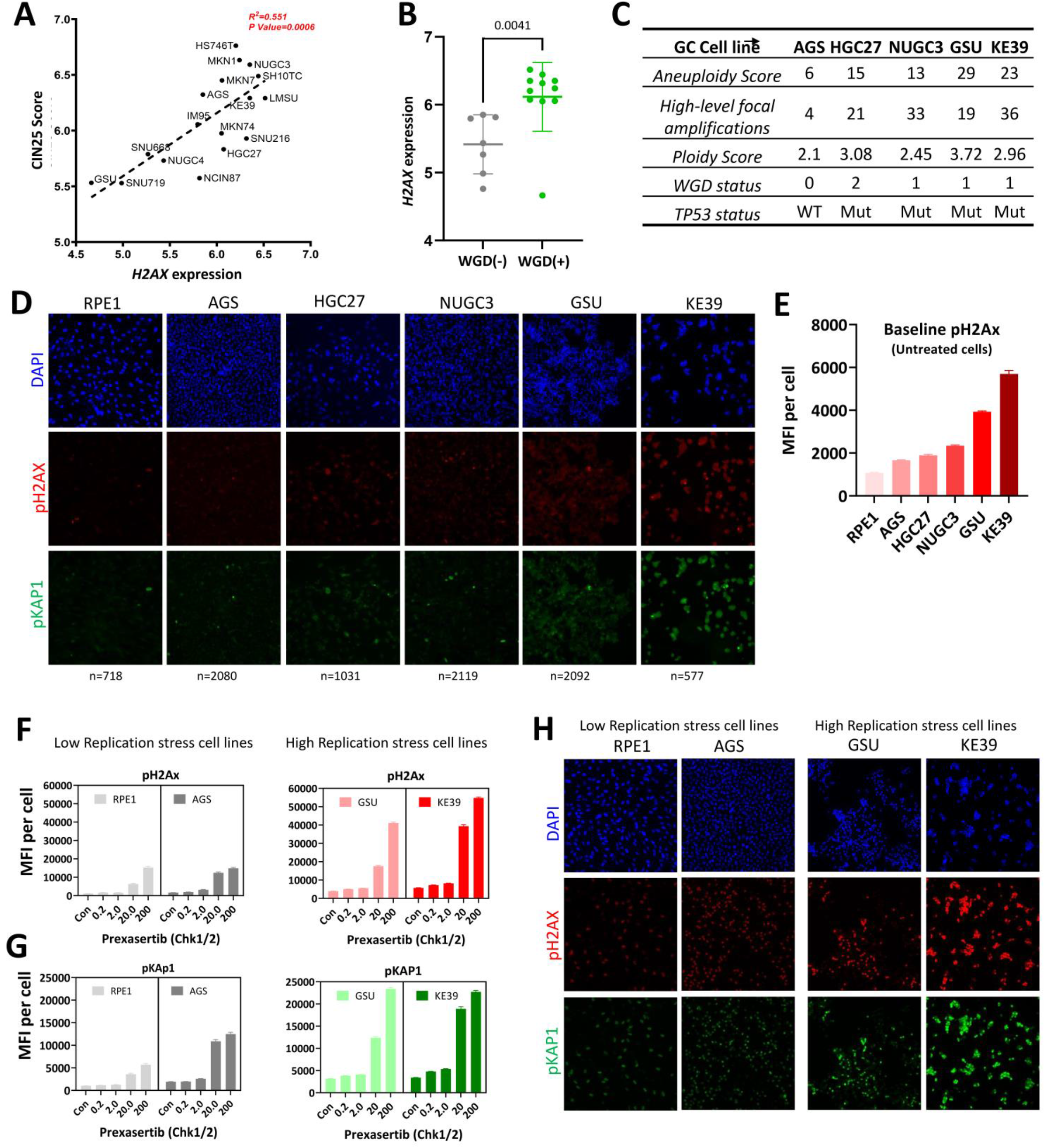
Aneuploidy correlates with DDR, H2AX expression, and replication stress in gastric cancer. A. Correlation plot between *H2AX* gene expression (log2(TPM+1)) and CIN25 score (DNA damage checkpoint expression) for the CCLE gastric cancer cell lines; R squared value and p-value calculated by simple linear correlation analysis. B. H2AX expression (log2(TPM+1)) between ‘No WGD (0)’ (grey) and ‘with WGD (1/2)’ (green) groups of gastric cancer cell lines available in the BROAD institute PRISM repurposing drug screen dataset; Difference between the H2AX expression is represented as the mean±S.D.; P-value calculated by unpaired t-test. C. Ploidy abnormalities scores for the five gastric cancer cell lines. D. Representative immunofluorescence images of pH2AX (red) and pKAP1 (green) stainings in untreated five gastric cancer cell lines and RPE1 cells (non-neoplastic cell line) representing intrinsic replication stress/double-stranded DNA breaks (DSBs); Nucleus stained with DAPI (blue). E. Quantification of data in D., expressed as mean fluorescence intensity per cell (MFI) ± S.D from the following number of cells-RPE1-718, AGS-2080, HGC27, 1031, NUGC3-2119, GSU-2092, KE39-577. F. Quantification of pH2AX staining in prexasertib (Chk1/2) dose-dependent induced replication stress in cell lines with ‘intrinsic low-replication stress’-RPE1, AGS (grey) and ‘intrinsic high-replication stress’-GSU, KE39 (red). Data expressed as mean fluorescence intensity (MFI) of pH2AX signal per cell ± S.D from the following number of cells-RPE1: Control-718, 0.2nM-888, 2.0nM-524, 20.0nM-570, 200.0nM-350; AGS: Control-2080, 0.2nM-1487, 2.0nM-1996, 20.0nM-1211, 200.0nM-1396; GSU: Control-2092, 0.2nM-3319, 2.0nM-4035, 20.0nM-2057, 200.0nM-2402; KE39: Control-577, 0.2nM-1004, 2.0nM-1022, 20.0nM-728, 200.0nM-958. G. Quantification of pKAP1 staining in prexasertib (Chk1/2) dose-dependent induced replication stress in cell lines with ‘intrinsic low-replication stress’-RPE1, AGS (grey) and ‘intrinsic high-replication stress’-GSU, KE39 (green). Data expressed as mean fluorescence intensity of pKAP1 signal per cell ± S.D from the following number of cells-RPE1: Control-718, 0.2nM-888, 2.0nM-524, 20.0nM-570, 200.0nM-350; AGS: Control-2080, 0.2nM-1487, 2.0nM-1996, 20.0nM-1211, 200.0nM-1396; GSU: Control-2092, 0.2nM-3319, 2.0nM-4035, 20.0nM-2057, 200.0nM-2402; KE39: Control-577, 0.2nM-1004, 2.0nM-1022, 20.0nM-728, 200.0nM-958. H. Representative immunofluorescence images of pH2AX (red) and pKAP1 (green) stainings in prexasertib (20nM) treated cell lines with intrinsic ‘low-replication stress’-RPE1, AGS, and intrinsic ‘high-replication stress’-GSU, KE39.

To validate the *in silico* analyses, we evaluated five gastric cancer cell lines (AGS, HGC27, HUGC3, GSU, and KE39) that showed varying levels of aneuploidy (Figure 3C) and one non-neoplastic diploid cell line (RPE1). We determined levels of pH2AX and pKAP1 in these cell lines to assess intrinsic DNA DSBs/replication stress. In response to DNA damage and replication stress, KAP1 is phosphorylated at serine 473(White et al., 2006). As expected, RPE1 cells showed minimal staining for pH2AX and pKAP1. Similarly, low levels of pH2AX and pKAP1 were observed in AGS, a TP53 WT gastric cancer cell line displaying the least aneuploidy defects. In contrast, we observed strong and moderate pH2AX and pKAP1 staining in high (KE39 and GSU) and intermediate (HGC27 and NUGC3) aneuploidy cell lines, respectively (Figure 3D). Importantly, since dsDNA breaks correlate with DDR response, these markers tracked with one another. As such, pH2AX and pKAP1 levels positively correlated with the degree of various ploidy defect measurements and p53 mutation status (Figure 3E). GSU and KE39 not only demonstrated greater baseline levels of DNA DSB and replication stress compared to RPE1 and AGS as measured by pH2AX and pKAP1, but also showed a more dramatic, dose-dependent induction when treated with prexasertib(Angius et al., 2020), a well-known Chk1/2 inhibitor (Figure 3F-H). These results demonstrate a correlation between broad measures of ploidy defects and DDR pathway activity in gastric cancer.

### Gastric cancer with high ploidy abnormalities and high *H2AX* expression are more sensitive to DDR pathway inhibitors

Based on these results, we hypothesized that, in addition to the degree of aneuploidy, the level of DDR pathway response to DNA DSBs and replication stress might correlate with sensitivity to drugs that inhibit DNA-damage checkpoint proteins. In other words, the effectiveness of DDR pathway inhibitors for gastric tumors may depend on levels of DDR activity in the setting of aneuploidy. To test this hypothesis, we used data from the PRISM repurposing drug screen of the BROAD Institute(Corsello et al., 2020) to characterize the sensitivity of the gastric cancer cell lines for DDR pathway inhibitors. Of all the inhibitors examined (Table S4), there was a positive association between higher aneuploidy score and sensitivity to multiple Chk1/2 inhibitors (prexasertib, PF-477736, and Rabusertib) (Figure 4A and Figure S4A). Moreover, higher aneuploidy score and Wee1 inhibitor MK1775 (AZD1775) sensitivity also showed a positive correlation (Figure 4G). Similarly, the CIN25 score in gastric cancer cell lines positively correlated with Chk1/2 inhibitor sensitivity (Figure S4B).

**Figure 4:**
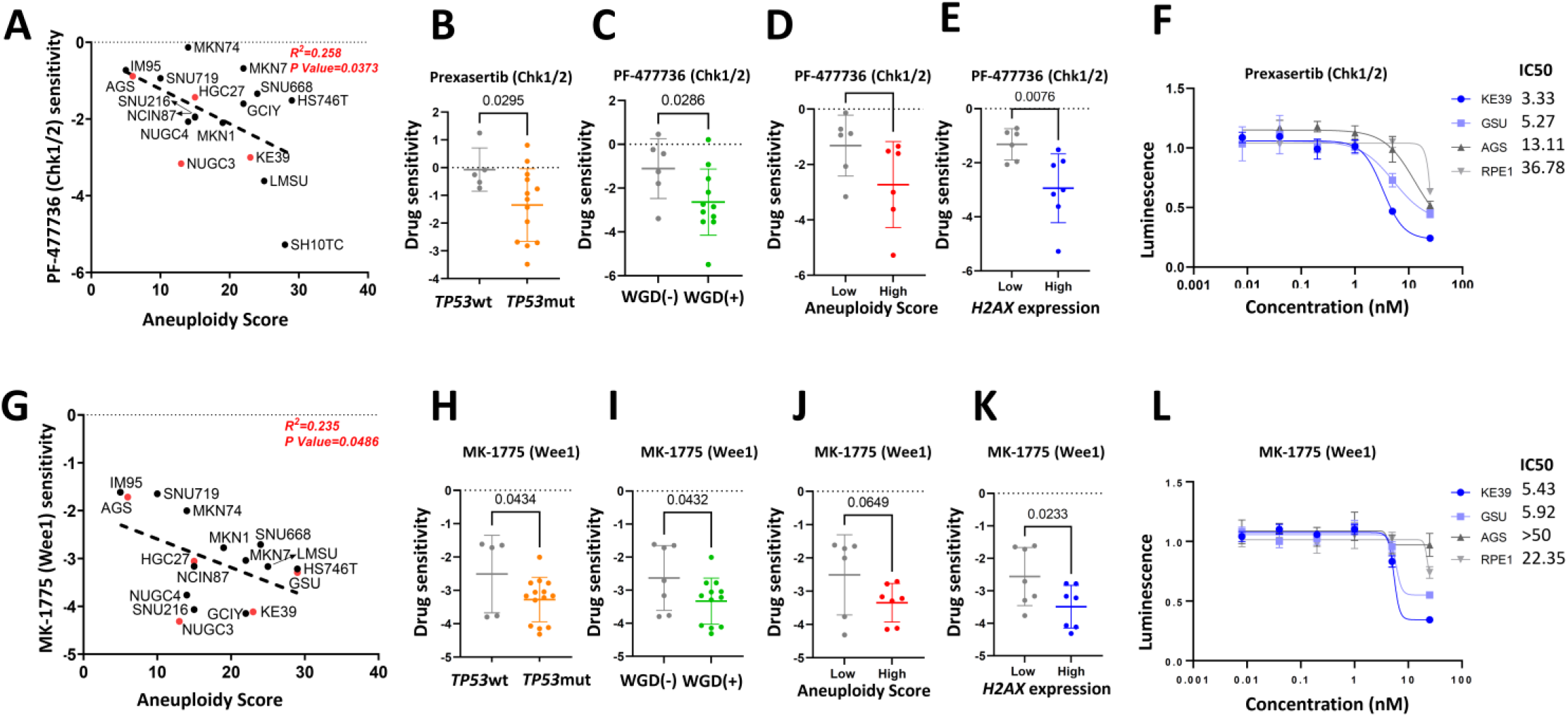
Gastric cancer with high ploidy abnormalities and elevated *H2AX* expression are more sensitive to DDR pathway inhibitors (Chk1/2 and Wee1) A. Correlation plot between aneuploidy score and PF-477736 (Chk1/2) sensitivity (log2 fold change) for the CCLE gastric cancer cell lines; Cell lines used in this study marked in red; R squared value and p-value calculated by simple linear correlation analysis. B. prexasertib (Chk1/2) sensitivity between ‘*TP53* wt’ and ‘*TP53* mut’ groups of gastric Cancer cell lines BROAD institute PRISM repurposing drug screen dataset; Difference between the prexasertib (Chk1/2) sensitivity (log2 fold change) is represented as the mean±S.D.; P-value calculated by unpaired t-test. C. PF-477736 (Chk1/2) sensitivity between ‘No WGD (0)’ (grey) and ‘with WGD (1/2)’ (green) groups of gastric cancer cell lines available in BROAD institute PRISM repurposing drug screen dataset; Difference between the PF-477736 (Chk1/2) sensitivity (log2 fold change) is represented as the mean±S.D.; P-value calculated by unpaired t-test. D. PF-477736 (Chk1/2) sensitivity between Low aneuploidy score (grey) and high aneuploidy score (red) groups of gastric cancer cell lines available in BROAD institute PRISM repurposing drug screen dataset; Difference between the PF-477736 (Chk1/2) sensitivity (log2 fold change) is represented as the mean±S.D.; P-value calculated by unpaired t-test. E. PF-477736 (Chk1/2) sensitivity between ‘low *H2AX* expressing’ (grey) and ‘high *H2AX* expressing’ (green) groups (log2(TPM+1)) of gastric cancer cell lines available in BROAD institute PRISM repurposing drug screen dataset; Difference between the PF-477736 (Chk1/2) sensitivity (log2 fold change) is represented as the mean±S.D.; P-value calculated by unpaired t-test. F. Dose-response curve of non-neoplastic cell line RPE1, Low replication-stress (grey)-AGS, and high replication-stress (blue)-KE39, GSU gastric cancer cell lines to indicated concentrations of prexasertib (Chk1/2); Best-fit IC50 scores are displayed; Data presented as mean±S.D. of four culture replicates at each indicated dose. G. Correlation plot between aneuploidy score and MK-1775 (Wee1) sensitivity (log2 fold change) for the CCLE gastric cancer cell lines; Cell lines used in this study marked in red; R squared value and p-value calculated by simple linear correlation analysis. H. MK-1775 (Wee1) sensitivity between ‘*TP53* wt’ and ‘*TP53* mut’ groups of gastric Cancer cell lines BROAD institute PRISM repurposing drug screen dataset; the difference between the MK-1775 (Wee1) sensitivity (log2 fold change) is represented as the mean±S.D.; P-value calculated by unpaired t-test. I. MK-1775 (Wee1) sensitivity between ‘No WGD (0)’ (grey) and ‘with WGD (1/2)’ (green) groups of gastric cancer cell lines available in BROAD institute PRISM repurposing drug screen dataset; Difference between the MK-1775 (Wee1) sensitivity (log2 fold change) is represented as the mean±S.D.; P-value calculated by unpaired t-test. J. MK-1775 (Wee1) sensitivity between low aneuploidy score (grey) and high aneuploidy score (red) groups of gastric cancer cell lines available in BROAD institute PRISM repurposing drug screen dataset; Difference between the MK-1775 (Wee1) sensitivity (log2 fold change) is represented as the mean±S.D.; P-value calculated by unpaired t-test. K. MK-1775 (Wee1) sensitivity between ‘low *H2AX* expressing’ (grey) and ‘high *H2AX* expressing’ (blue) groups (log2(TPM+1)) of gastric cancer cell lines available in BROAD institute PRISM repurposing drug screen dataset; Difference between the MK-1775 (Wee1) sensitivity (log2 fold change) is represented as the mean±S.D.; P-value calculated by unpaired t-test. L. Dose-response curve of non-neoplastic cell line RPE1, Low replication-stress (grey)-AGS, and high replication-stress (blue)-KE39, GSU gastric cancer cell lines to indicated concentrations of MK-1775 (Wee1); Best-fit IC50 scores are displayed; Data presented as mean±S.D. of four culture replicates at each indicated dose.

Based on these observations and to further build confidence in our results, we divided the gastric cancer cell lines into 2 groups (quartiles) based on *TP53* status and ‘high or low’ ploidy abnormalities (aneuploidy score, WGD, CIN25 score, high amplitude focal amplifications, and ploidy score). Gastric cancer cell lines with *TP53* mutation and a high degree of ploidy abnormalities were significantly more sensitive to the Chk1/2 (Figure 4B-D) and Wee1 (Figure 4H-J) inhibitors (S4C-G). We also observed that gastric cancer cell lines with specific ploidy defects were sensitive to a subset of ATR inhibitors (AZD6738 for high ploidy cell lines and ETP-46464 for high CIN25 cell lines), but validation experiments using AZD6738 were not able to confirm these results (Figure S4H).

The expression of *H2AX* in gastric cancer cell lines also positively correlated with Chk1/2 inhibitor sensitivity (Figure S4I). Gastric cancer cell lines with high *H2AX* expression were also more sensitive to Chk1/2 and Wee1 inhibitors (Figure 4E, K and S4J). In contrast, gastric cancer cell lines with high ploidy abnormalities were not sensitive to chemotherapeutic agents routinely used in the clinic to treat patients with gastrointestinal cancers. As an example, we show the sensitivity of gastric cancer cell lines with high and low WGD to 8 chemotherapeutic agents (Figure S4K). DDR inhibitors that did not show a significant difference among gastric cancer cell lines with high or low ploidy abnormalities are summarized in Figure S4L-N; there is an enrichment of ATR and ATM inhibitors among these agents.

Encouraged by these results, we validated that gastric cancer cell lines with high ploidy defects (GSU/KE39) showed greater sensitivity to Chk1/2 and Wee1 inhibitors compared to those with low ploidy defects (RPE1/AGS) (Figure 4F, L). These results are in agreement with the correlation plot between aneuploidy and DDR inhibitor sensitivity (Figures 4A and 4G), in which the cell lines used for validation are marked in red. Collectively, these results suggest that biomarkers of DNA DSBs/replication stress and ploidy defects could be predictive markers for DDR pathway inhibitor sensitivity in gastric cancer.

### Cumulative score of ploidy abnormalities (CSPA) predicts prexasertib (Chk1/2) response

Given the predictive value of individual measures of aneuploidy and DDR pathway activity, we asked whether we could combine these parameters to better predict sensitivity to DDR inhibitors. Indeed, when we combined these parameters using a simple scoring scheme, which we call the cumulative score of ploidy abnormalities (CSPA), we improved the correlation with prexasertib sensitivity (R^2^=0.623, P=0.0002) compared to individual values (*TP53* status, ploidy abnormalities, and *H2AX* expression) (Figure 5A and S5A). Moreover, binning the cell lines into low, medium, and high CSPA yielded a significant difference for prexasertib sensitivity, with high CSPA cell lines showing the greatest prexasertib (Chk1/2 inhibitor) sensitivity (Figure 5B and S5B). As expected, CSPA showed a strong positive correlation with DNA-PKcs (*PRKDC*), a specific marker of DDR activity (Figure S5C). The results from our cumulative score approach suggest that instead of using only one parameter, a combined score, such as CSPA, is a better predictor for DDR inhibitor sensitivity.

**Figure 5:**
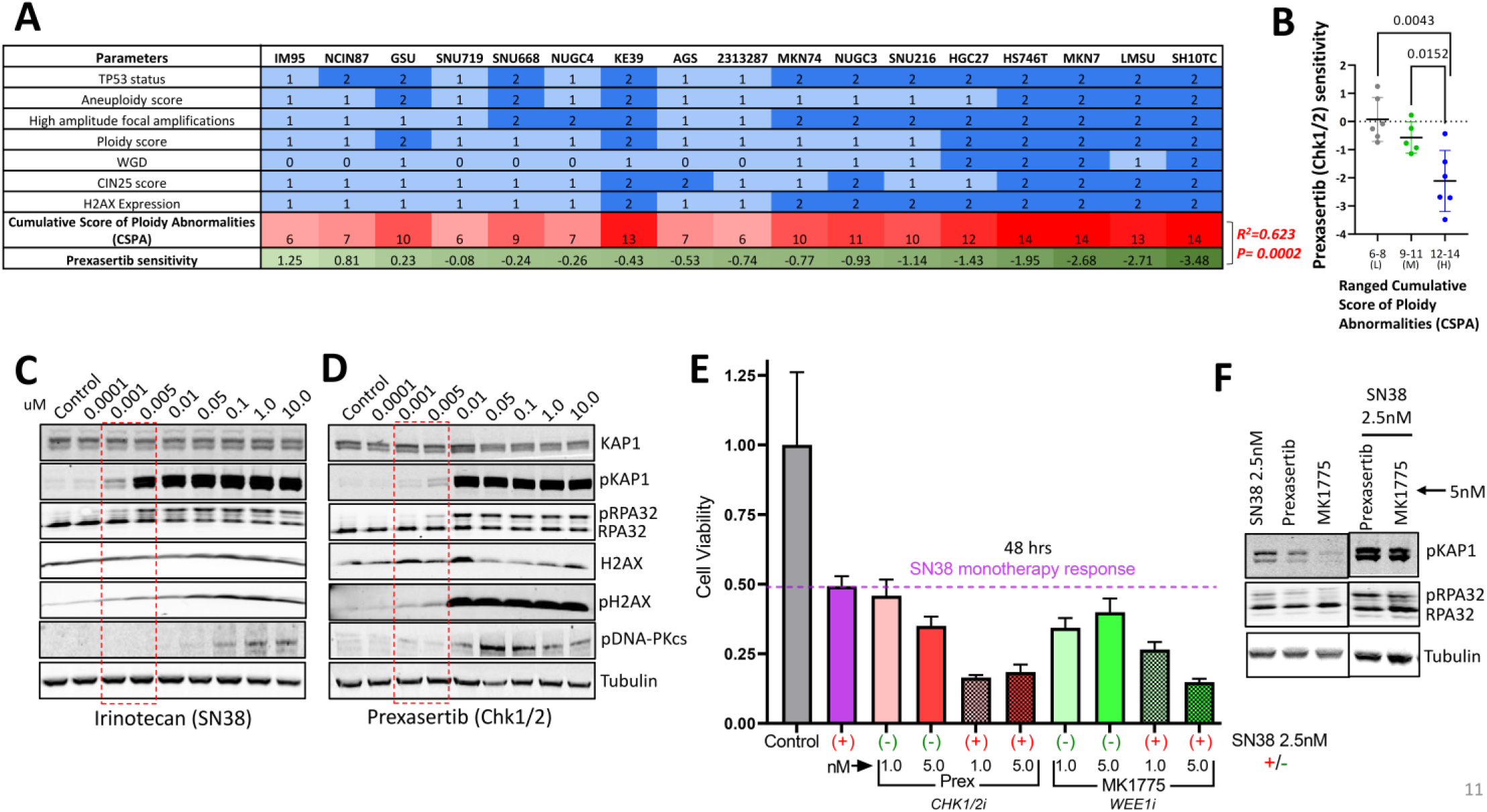
Cumulative score of ploidy abnormalities predicts DDR pathway inhibitor response in gastric cancer cell lines. A. Heatmap table representing the correlation between the Cumulative score of ploidy abnormalities and prexasertib sensitivity for the CCLE gastric cancer cell lines. The scoring scheme is as follows: TP53 status: wt=1, mut=2; Aneuploidy score: 15 and less than 15=1, more than 15=2; High amplitude focal amplifications: 20 and less than 20=1, more than 20=2; Ploidy score: 2.5 and less than 2.5=1, more than 2.5=2; WGD: No WGD=0, WGD 1=1, WGD2=2; CIN25 score: 6.1 and less than 6.1=1, more than 6.1=2; H2AX Expression: 6 and less than 6=1, more than 6=2. R squared value and p-value calculated by simple linear correlation analysis. B. prexasertib (Chk1/2) sensitivity between three sub-ranged ‘Cumulative score of ploidy abnormalities’ groups of gastric Cancer cell lines BROAD institute PRISM repurposing drug screen dataset; Difference between the prexasertib (Chk1/2) sensitivity (log2 fold change) is represented as the mean±S.D.; L=low, M=medium, H=high; P-value calculated by unpaired t-test. C. Immunoblot of replication stress/DDR markers and tubulin as the loading control in the KE39 cell line (high intrinsic replication stress) treated with the indicated concentrations of irinotecan (SN38). D. Immunoblot of replication stress/DDR markers and tubulin as the loading control in the KE39 cell line (high intrinsic replication stress) treated with the indicated concentrations of prexasertib (Chk1/2). E. Proliferation of the KE39 cell line (high intrinsic replication stress) treated with vehicle (grey), 2.5 nM of SN38 (magenta), prexasertib (red), prexasertib+SN38 (pattered red), MK1775 (green), MK1775+SN38 (pattered green); Light shade of color-1.0nM of DDR inhibitor; Dark shade of color-5.0nM of DDR inhibitor; All treatments for 48 hrs. F. Immunoblot of pKAP1, pRPA32, and tubulin as the loading control in KE39 cell line (high intrinsic replication stress) treated with monotherapy-SN38 (2.5nm), prexasertib (5nM), MK1775 (5nM) or combination therapy-SN38 (2.5nM)+prexasertib (5nM) and SN38 (2.5nm)+MK1775 (5nM).

To understand the relationship between DDR pathway and chemotherapy response in patients with gastric cancer, we decided to examine previously established Recombination Proficiency Score (RPS)(Pitroda et al., 2014) in human gastric cancer gene expression data. RPS can predict adverse clinical features and response to chemotherapy, especially topoisomerase-I inhibiting drugs like topotecan and irinotecan, the latter of which has routinely been used in GEA. Lower RPS depicts inferior patient survival rates, higher genomic stability, and more sensitivity to chemotherapy. In line with our TCGA data analysis, mutant *TP53* gastric cancer in TCGA have lower RPS than wild-type *TP53* patients (Figure S5D), confirming that patients with mutant *TP53* gastric cancer have a poor prognosis and higher sensitivity to irinotecan. Moreover, *H2AX* expression correlates negatively with RPS score in human gastric cancer. Patients with low RPS gastric cancers display higher *H2AX* expression, reinforcing our confidence in *H2AX* expression as a biomarker for tumors exhibiting higher DNA damage and better therapy response (Figure S5E).

Chemotherapy is the backbone of most treatment regimens for gastric cancer. We hypothesized that gastric cancers with high DNA DSBs/replication stress would be more sensitive to chemotherapy and DDR pathway inhibitors combination. We therefore sought to identify chemotherapy-DDR pathway inhibitors combinations that enhance cytotoxocity, with the goal of nominating potential clinical trial concepts. KE39 cell line exhibited high DDR pathway activity and therefore a suitable model for these experiments. Based on the recent preclinical and clinical studies, gemcitabine, a chemotherapeutic agent that generates replication stress, showed evidence of synergy when combined with DDR pathway inhibitors in small cell lung cancer, pancreatic cancer, and ovarian cancer(Konstantinopoulos et al., 2020; Sen et al., 2019; Wallez et al., 2018). Irinotecan is also a commonly used chemotherapeutic in the treatment of gastrointestinal cancers. We therefore examined the combination of either gemcitabine or irinotecan with Chk1/2 or Wee1 DDR pathway inhibitors. It is worth noting that while irinotecan is used clinically in gastroesophageal cancer, gemcitabine is not. KE39 cells were generally more sensitive to SN38, the active metabolite of irinotecan(Ramesh et al., 2010) (Figure S5F), than gemcitabine-treated cells. Consistently, SN38 elicited activation of DDR pathway as measured by H2AX, pH2AX, and pDNA-PKcs as well as replication stress as measured by pKAP1 and pRPA32 at lower doses than gemcitabine (Figure 5C and S5G, effect at 1nM for SN38 compared to 10nM for gemcitabine). For the DDR pathway inhibitor prexasertib, the increase in DDR/replication stress marker was first observed at 1nM, with a more substantial increase at 5nM (Figure 5D). When SN38 (2.5nM) was combined with prexasertib or Wee1 inhibitor MK1774 (1.0nm and 5.0nm), the combinations showed a better effect in reducing cell viability and inducing DDR/replication stress compared with the individual monotherapies (Figure 5E-F and Figure S5H). We did not observe this synergy when we combined prexasertib or MK1775 (Wee1) with gemcitabine (Figure S5I).

Earlier in this study, we observed an increase in p21 staining during the premalignant to malignant progression in human gastric cancer specimens and in lesions from MNU-treated mice, and found that this staining pattern is independent of p53 status. For SN38 treatment in KE39 cells, the dose-dependent induction in DDR/replication stress correlates with an increase in p21 expression (Figure S5J). This observation is particularly important because KE39 cells harbor a loss-of-function *TP53* mutation. In contrast to the combination with SN38, the combination of gemcitabine with prexasertib or MK1775 was not potent enough to increase DDR/replication stress or p21 levels (Figure S5K). Cumulatively, these clinically relevant data suggest that the combination of irinotecan with Chk1/2 or Wee1 inhibitors is a promising treatment strategy for patients with gastric cancers that display elevated biomarkers of DDR.

## DISCUSSION

CIN constitutes the largest subtype of gastric cancer, and most of these tumors are heterogeneous, highly aneuploid, and harbor loss-of-function *TP53* mutations (Giam and Rancati, 2015). Replication stress is common cellular stress associated with aneuploidy that results from CIN (Zhu et al., 2018). Moreover, aneuploidy generates replication stress-mediated DNA DSBs, activating the DDR pathway (Ohashi et al., 2015). Failure of cell-cycle control can lead to WGD(Dewhurst et al., 2014; Goyal et al., 2011; Quinton et al., 2021a). Cancer cells with high replication stress are subjected to ploidy abnormalities such greater focal amplifications. As such, CIN gastric cancer is intrinsically challenged by adaptive responses to aneuploidy, such as DDR, which may also reveal unique vulnerabilities that can be exploited for cancer therapy (Quinton et al., 2021b; Zhu et al., 2018). The first part of our study aimed to better understand the contribution of DDR/replication stress in premalignant progression of human gastric cancer.

We determined that gastric premalignancy is associated with increased expression of specific DDR markers using matched normal, premalignant, and malignant lesions from 8 patients with gastric cancer and a carcinogen-induced mouse model. The cell cycle markers p53, p16, and p21 showed striking heterogeneity within the same tumor and between the precancerous/cancerous lesions from different samples. For both the patients and mouse model, p21 gradually increased from normal to premalignant to malignant lesions independent of p53 status. p21 is downstream of p53, and its expression independent of p53 has been previously reported in other cancers (Origanti et al., 2013). However, the precise mechanism of how p21 becomes activated in the absence of p53 requires further study. A possible mechanism may be that regulatory feedback systems upregulate p21 in the absence of p53.

Our findings reveal significantly higher expression of DDR markers *H2AX* and *TP53BP1* in stomach adenocarcinomas compared with normal tissue. We also found that higher *H2AX* expression is found in *TP53* mutant gastric cancer, and that *TP53* mutations are predominant in the intestinal and CIN subtype of gastric adenocarcinoma. H2AX is a marker of activated DNA damage and has been previously implicated in cancer progression (Bartkova et al., 2005; Krishnan et al., 2020). In addition, we observed that higher expression of *H2AX* correlates with CIN and WGD in gastric cancer. Furthermore, patients with gastric adenocarcinoma that exhibit low RPS indicative of elevated genome instability, adverse clinical features, and enhanced sensitivity to chemotherapy, especially irinotecan, have higher *H2AX* expression. These findings demonstrate that DNA damage and CIN are strongly associated in the evolution of gastric cancer, and that elevated markers of DDR may predict response to treatment.

In line with these results in patients, we found that *TP53* mutation, ploidy abnormalities, and H2AX expression independently positively correlate with sensitivity to Chk1/2 and Wee1/2 DDR pathway inhibitors in gastric cancer cell models. The predictive power for DDR pathway sensitivity was improved when these parameters of aneuploidy and DDR were combined. These findings suggest that ploidy abnormalities and *H2AX* expression may have clinical value as biomarkers to predict response to DDR pathway inhibitors (Chk1/2 and Wee1) in patients with gastric and esophageal adenocarcinoma. We further these findings by demonstrating promising data that combining irinotecan with DDR pathway inhibitors improves anti-tumor response in a model of gastric cancer. We hope our study will open new avenues for utilizing DDR markers and ploidy defects as diagnostic proxy that may predict premalignant progression and as an indication for using DDR pathway inhibitors in the treatment of a subset of patients with gastric and esophageal adenocarcinoma.

## LIMITATIONS OF THE STUDY

It was challenging to find patient specimen with paired premalignant and malignant lesions. We therefore had limited samples to analyze (n = 8), reducing the power of our histopathological analysis. Our in vitro studies were limited to 5 gastric cancer cell lines and one non-neoplastic cell line. Further experimentation in organodis and patient-derived xenografts (PDXs) are warranted to improve preclinical data. We used prexasertib and MK-1775 as potent inhibitors of chk1/2 and Wee1, respectively, as these DDR inhibitors are commonly used in clinical trials for solid tumors. There are however next-generation DDR inhibitors that may prove less toxic and more efficacious; our results can therefore be used as a proof-of-concept.

## AUTHOR CONTRIBUTIONS

Conceptualization: P.S., N.S.S.; Methodology: P.S., D.P., Z.M.S.; Validation: P.S., D.P., Z.M.S.; Formal Analysis: P.S., D.P., Z.M.S.; Investigation: P.S., N.S.S.; Data Curation: P.S., Z.M.S., A.P.; Writing – Original Draft: P.S.; Writing: Review & Editing: P.S., D.P., Z.M.S., V.T., S.S., B.H., A.P., H.S., J-B.Z., Z.S., J.M.C, N.S.S.; Visualization: P.S., D.P., Z.S.; Supervision: Z.S., N.S.S.; Funding Acquisition: P.S., N.S.S.

## ACKNOWLEDGMENTS

The authors thank the pathology core facility of Brigham and Women’s Hospital and Beth Israel Deaconess Medical Center for assistance with histopathology; Haley Elizabeth Szewczuga at DFCI for help in editing the manuscript; members of the Sethi lab for insightful discussions. P.S. is Funded by the Finnish cultural foundation and Sigrid Juselius foundation. This study is funded by philanthropic support from Pan-Mass Challenge Team Detect-Me-If-You-Can (Nick Tierney Gastric Cancer Fund) to the Sethi Lab, the National Institute of Health NIDDK K08 (DK120930) (N.S.S), Augustyn Award in Digestive Cancer from the American Gastroenterology Association (N.S.S), Gastric Cancer Foundation (N.S.S), and a DeGregorio Family Foundation Award (N.S.S).

## MATERIALS AND METHODS

### Histopathology and Immunohistochemistry

For immunostaining, antigen retrieval was performed using a sodium citrate buffer (pH6), Trilogy (Sigma Aldrich Cell Marque), or Tris-EDTA pH9. Slides were permeabilized using a 0.2% Triton X100 for 30 minutes at room temperature and blocked with donkey serum for 1 hour. The primary antibodies used for immunohistochemistry were: Human tissue-rabbit anti-ki67 (1:1000, Biocare Medical, #CRM325-5b6), rabbit anti-H2AX (1:1000, CST, #7631), rabbit anti-pH2AX ser139 (1:1000, CST, #9718), mouse anti-53BP1 (1:1000, Millipore, #3802), mouse anti-p53 (1:1000, CST, #48818), mouse anti-P16INK4 (1:1000, Abcam, #54210), rabbit anti-p21 (1:1000, CST, #2947), Murine tissue-rabbit anti-ki67 (1:1000, Biocare Medical, #CRM325-5b6), rabbit anti-pH2AX ser139, (1:1000, CST, #9718), rabbit anti-p53 (1:1000, Abcam, #1431), rabbit anti-p21 (1:1000, Abcam, #188224). Binding of primary antibody was detected with 3,39-diamino-benzidine-tetrahydrochloride-dihydrate and counterstained with hematoxylin. Images were acquired with Leica DM750 microscope. The slides were reviewed by expert GI pathologist (D.T.P.). For each antibody, the percentage positive cells (<=5%, 6-50%, and >50%) were assessed along with intensity of staining (weak, moderate, or strong). For p53, overexpression and null pattern of expression were considered “abnormal” expression while weak, patchy staining was considered normal/wild-type expression. Cytoplasmic and nuclear p16 expression were assessed in all cases.

### Cell lines and cell culture

Human gastric cancer cell lines were obtained from the CCLE core facility (BROAD Institute, Cambridge), which obtained them directly from commercial sources and authenticated the lines using standard short tandem repeat analysis. RPE1 cells were obtained from the American Type Culture Collection (ATCC). RPE1 cells were grown in DMEM-F12 (Life Technologies, #10565042) supplemented with 10% FBS and 1% penicillin/streptomycin. HGC27 cells were grown in DMEM (Life Technologies, #11965118) supplemented with 10% FBS and 1% penicillin/streptomycin. AGS, GSU, KE39, and NUGC3 were grown in RPMI 1640 (Life Technologies, #11875119) supplemented with 10% FBS and 1% penicillin/streptomycin. Cells lines were maintained in a humidified 37°C incubator with 5% CO2 and routinely tested for mycoplasma contamination (Lonza #LT07-118).

### Immunofluorescence

Cells were plated on glass-Teflon microscope slide (Tekdon, #518plain) and the next day treated with either DMSO or different concentrations of prexasertib (Selleck, #S7178) for 24 hours. Cells were then washed and fixed with 4% paraformaldehyde (PFA) for 15 minutes at room temperature, blocked and permeabilized PBS-BSA + 0.3% Triton X-100 for 15 minutes at room temperature, and incubated with primary antibodies (rabbit anti-pTIF1β/pKAP1 (Ser824) (1:500, CST # 4127S) and Mouse anti-pH2AX (1:600, CST #80312) in PBS-BSA overnight at +4°C. After 3 washes with PBS, secondary antibodies; anti-Rabbit IgG Alexa Fluor 488 (Life tech #A11008) and anti-Mouse IgG Alexa Fluor 568 (Life tech #A11004), diluted 1:300 in PBS-BSA with DAPI (1:1000) for 2 hours at room temperature. Finally, the cells were washed with PBS 3 times, mounted, and covered in aluminum foil for imaging.

### Microscopy and Image quantification

Immunofluorescence imaging was performed using Nikon Eclipse Ti2 Series inverted microscope (×40 objective). NIS-Elements AR software was used to acquire the images. DAPI channel was used to focus the cells, and the images for 488 (pKAP1) and 567 (pH2AX) channels were acquired, keeping the same exposure across the samples. For image analysis, the 16-bit images were converted to 2-bit images in ImageJ, followed by ‘hole filling’ and segmentation (watershed) commands. The cells were then automatically recognized and counted using ‘analyze particles’, and the file was saved as an ‘image mask’. Finally, the image mask was overlaid on the original green (488) and red (567) split channels, and the ‘measure’ command was used to get the mean fluorescence intensity from each cell.

### Cell Proliferation Assay and inhibitors

For IC50 experiments, 1000 cells were plated in a flat-bottom 96-well plate. Cells were treated with either vehicle (DMSO) or different concentrations of the inhibitors. Luminescence was measured using CellTiter-Glo (Promega, #G7572) for ATP amount after 3 days, and final readings were normalized with Day 1 luminescence readings. For colony formation assay, 2×10^4^-1×10^5 cells were plated in 6-well plates. Cells were then treated with DMSO or inhibitors. After 4-6 days, cells were fixed in 1% paraformaldehyde for 15 minutes at room temperature, washed twice with PBS, and stained with 0.1% crystal violet solution in ethanol (Sigma Aldrich, #HT901) for 15 minutes at room temperature. ImageJ was used to quantify the mean intensity of the scanned plates. The following drug agents were used in the study: prexasertib (Selleck, #S7178), AZD6738 (Selleck, #S7693), MK1775 (Selleck, #S1525), gemcitabine (Selleck, #S1149), and SN-38 (Selleck, #S4908).

### Immunoblot

Cells were washed with cold phosphate buffer saline (PBS) and lysed in RIPA buffer (Sigma-Aldrich, #R0278-50ML) supplemented with a protease inhibitor (Roche, #11873580001) and phosphatase inhibitor (CST #5870S). Protein extracts were separated using SDS-PAGE under denaturing conditions (8–16% Novex™ Wedgewell™ Tris-Glycine Gels) and were transferred to PVDF membranes (iBlot2™ PVDF Regular Stacks). Membranes were blocked with blocking buffer (InterceptTM Blocking buffer, #927-60001) and incubated with the primary antibodies diluted in blocking buffer and incubated overnight at +4°C. After primary antibody incubation: rabbit anti-pTIF1β/pKAP1 (Ser824) (1:1000, CST # 4127S), rabbit anti-TIF1β/KAP1 (1:1000, CST # 4124S), rabbit anti-H2AX (1:1000, CST #7631), mouse anti-pH2AX (1:1000, CST #80312), rabbit anti-pDNA-PKcs (1:1000, CST, #12311S), rabbit anti-αTubulin (1:5000, CST, #2125), rabbit anti-p21 (1:500, CST, #2947S), rabbit anti-RPA32 (1:1000, Bethyl, A300-244A) and rabbit anti-pRPA32 (1:1000, Bethyl, A300-246A) membranes were washed 3 times with TBST and incubated with fluorophore-conjugated secondary antibodies diluted (1:10,000) in blocking buffer at room temperature for 1 hour. Membranes were then scanned using an infrared imaging system (Odyssey; LI-COR Biosciences). The following secondary antibodies were used: donkey anti-mouse IRDye 800CW (LI-COR, 926-32212), donkey anti-mouse IRDye 680RD (LI-COR, 926-68072), donkey anti-rabbit IRDye 800CW (LI-COR, 926-32213), and donkey anti-rabbit IRDye 680RD (LI-COR, 926-68073).

### TCGA analysis

For the TCGA dataset FPKM-UQ, normalized expression values were downloaded from the TCGA data portal and subtype information was accessed using TCGABiolinks R package(Colaprico et al., 2016). The following subtypes were defined: CIN (chromosomal instability), GS (genomically stable), EBV (Epstein–Barr virus-positive), MSI (microsatellite instability), and HM-SNV (hypermutated-single-nucleotide variant predominant), based on(Liu et al., 2018). World Health Organization (WHO) histological classification was available for these samples and Lauren-classification was determined based on WHO classification. *TP53* mutation status was downloaded from cBioPortal and only pathogenic mutations were treated as mutant cases.

### Drug sensitivity and correlation analysis

The gene expression, aneuploidy, and drug sensitivity data were downloaded (22Q2 data release from DepMap website of Broad Institute [https://depmap.org]). For the drug sensitivity analysis, the cell lines were divided into a high or low group based on quartiles of the expression of gene of interest, score, or feature. The difference between the mean of sensitivity for the two groups was plotted, and the p-value was calculated using the unpaired t-test. For the correlation analysis, the continuous data were plotted, and R-squared/p-value was calculated using simple linear correlation analysis.

