## Supplementary Files for "Replicative stress in gastroesophageal adenocarcinoma is associated with chromosomal instability and sensitivity to DNA damage response inhibitors"

### SUPPLEMENTAL TABLES

- **Table S1:** Confirmation of the normal, premalignant, and malignant status of gastric tissue in each patient based on Ki-67 staining.
- **Table S2**: Patient-wise summary of staining pattern of select DDR markers (53BP1, H2AX, and pH2AX) in paired premalignant and malignant gastric lesions.
- **Table S3:** Patient-wise summary of staining pattern of cell cycle regulation markers in paired normal, premalignant, and malignant gastric lesions.
- **Table S4:** List of the DDR pathway inhibitors analyzed in this study and their targets

**SUPPLEMENTAL FIGURES AND LEGENDS**

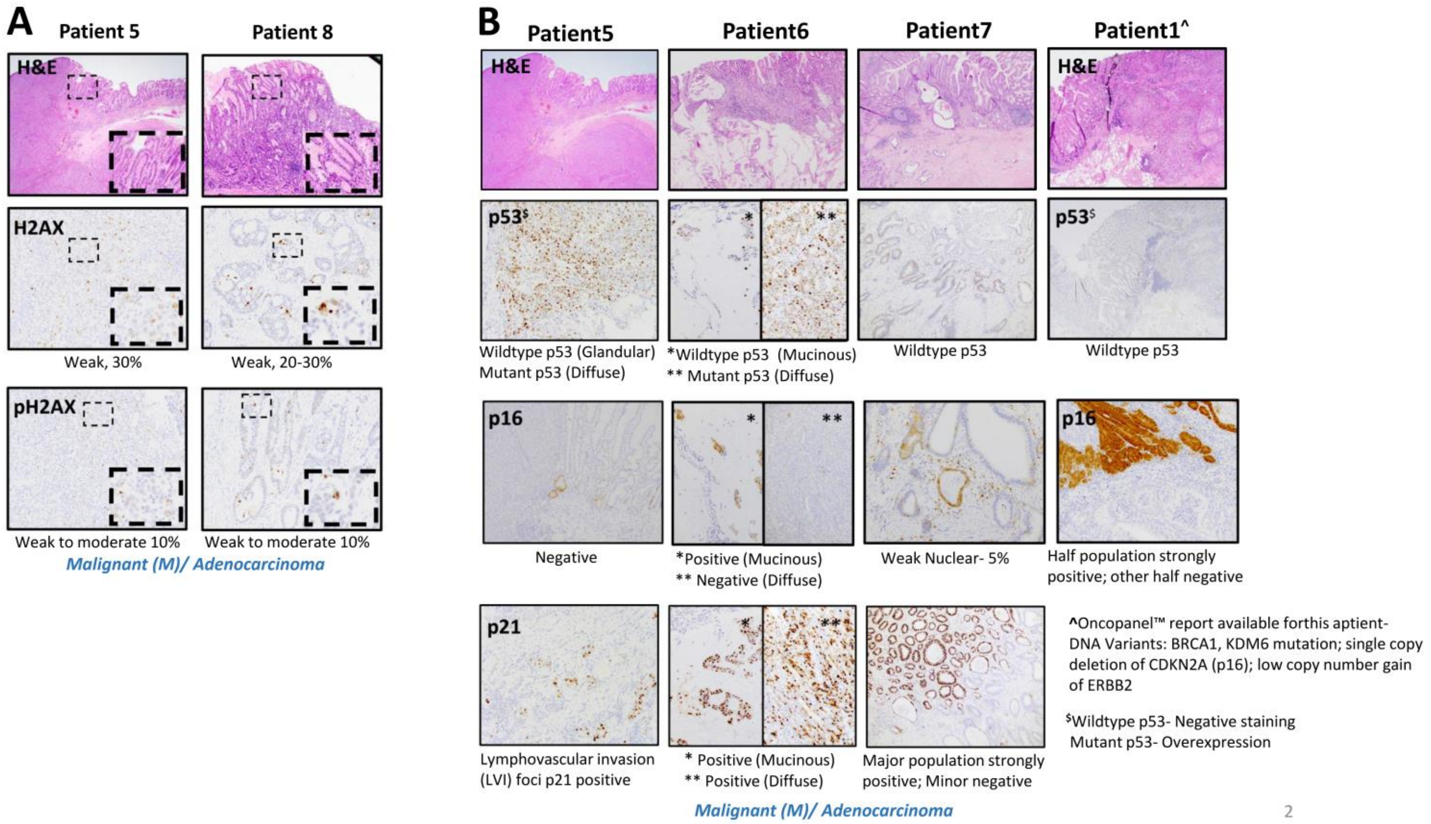


**Figure S1: Staining pattern of DDR marker H2AX/pH2AX and Cell cycle regulation markers in human gastric adenocarcinoma**

A. IHC images of patient5 and patient8 displaying weak to moderate H2AX and pH2AX levels in the malignant gastric lesion. Insets represent the zoomed area for each image.
B. Representative IHC images of p53, p16, and p21 stainings in malignant lesions from four patients representing intratumor and intertumor heterogeneity in staining patterns.

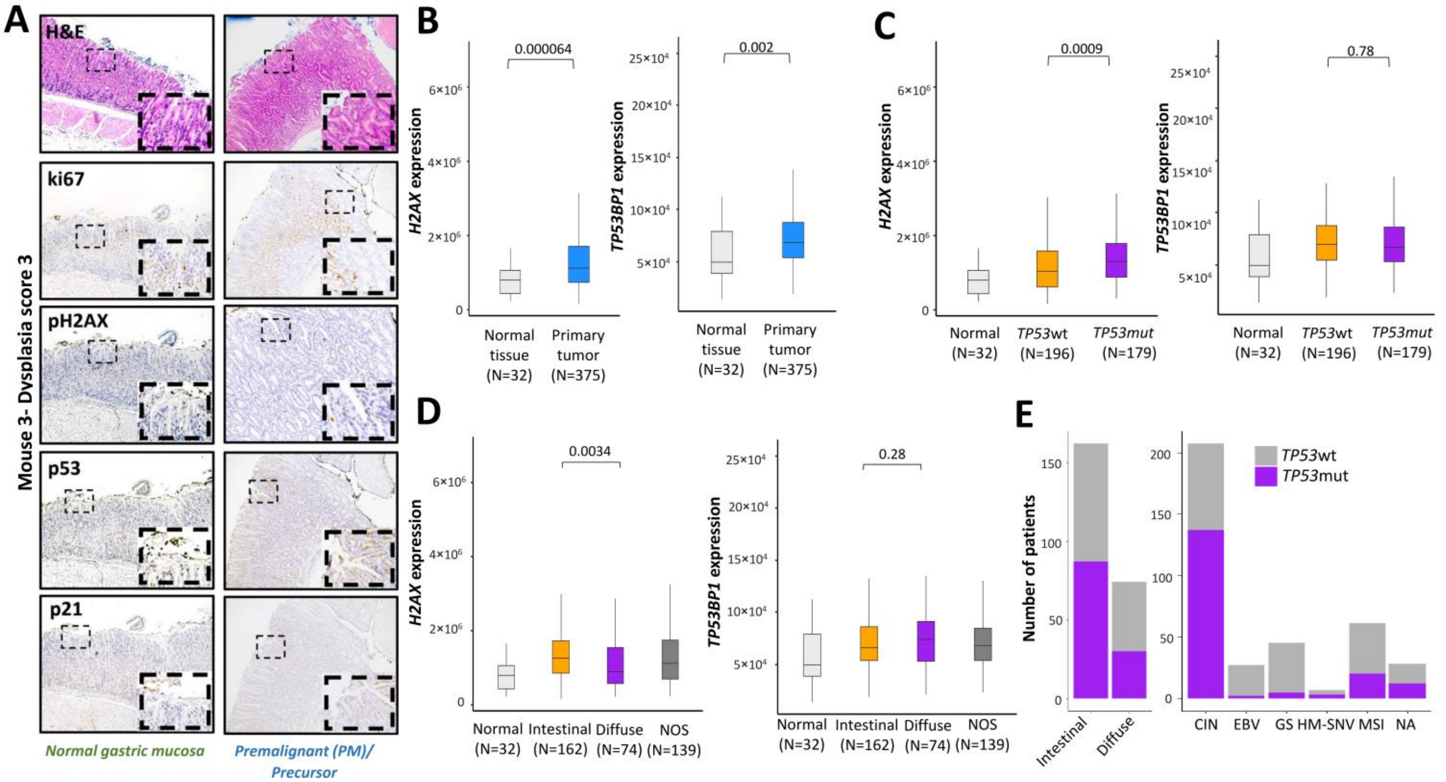

**Figure S2: Evaluation of DDR markers/cell cycle regulation markers in a carcinogen-induced mouse model of gastric premalignancy and *H2AX/TP53BP1* expression in TCGA gastric adenocarcinoma patients (STAD).**

A. Representative IHC images of staining pattern of proliferation marker ki67, DDR marker pH2AX, and cell cycle regulation markers (p53 and p21) in paired normal and premalignant gastric lesions of MNU treated mouse3 (dysplasia score 3). p53 and p16 show heterogeneous and inverse staining pattern.
B. Expression (FPKM) of *H2AX* and *TP53BP1* in gastric normal tissue and tumor tissue from the Stomach adenocarcinoma (STAD) cohort of TCGA.
C. Expression (FPKM) of *H2AX* and *TP53BP1* in gastric normal tissue, '*TP53*wt' or '*TP53*mut' tumors from Stomach adenocarcinoma (STAD) cohort of TCGA.
D. Expression (FPKM) of *H2AX* and *TP53BP1* in Lauren classification subtypes of gastric cancer from Stomach adenocarcinoma (STAD) cohort of TCGA.
E. Number of patients with '*TP53*wt' or '*TP53*mut' status in Lauren classification and molecular classification subtypes of gastric cancer patients from Stomach adenocarcinoma (STAD) cohort of TCGA.

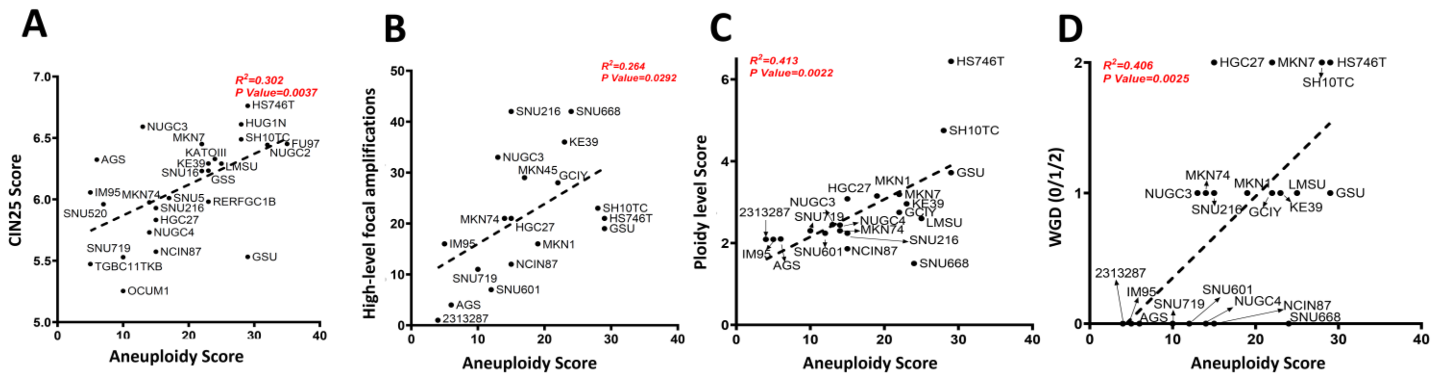


**Figure S3: Aneuploidy in gastric cancer correlates with other ploidy abnormalities in gastric cancer (high-level focal amplification, ploidy, and WGD)**

A. Correlation plot between aneuploidy score and CIN25 score (DNA damage checkpoint expression) for the CCLE gastric cancer cell lines. R squared value and p-value calculated by simple linear correlation analysis.

B. Correlation plot between aneuploidy score and number of high-level focal amplifications score for the CCLE gastric cancer cell lines. R squared value and p-value calculated by simple linear correlation analysis.

C. Correlation plot between aneuploidy score and ploidy level score for the CCLE gastric cancer cell lines; R squared value and p-value calculated by simple linear correlation analysis.

D. Correlation plot between aneuploidy score and WGD for the CCLE gastric cancer cell lines; R squared value and p-value calculated by simple linear correlation analysis.

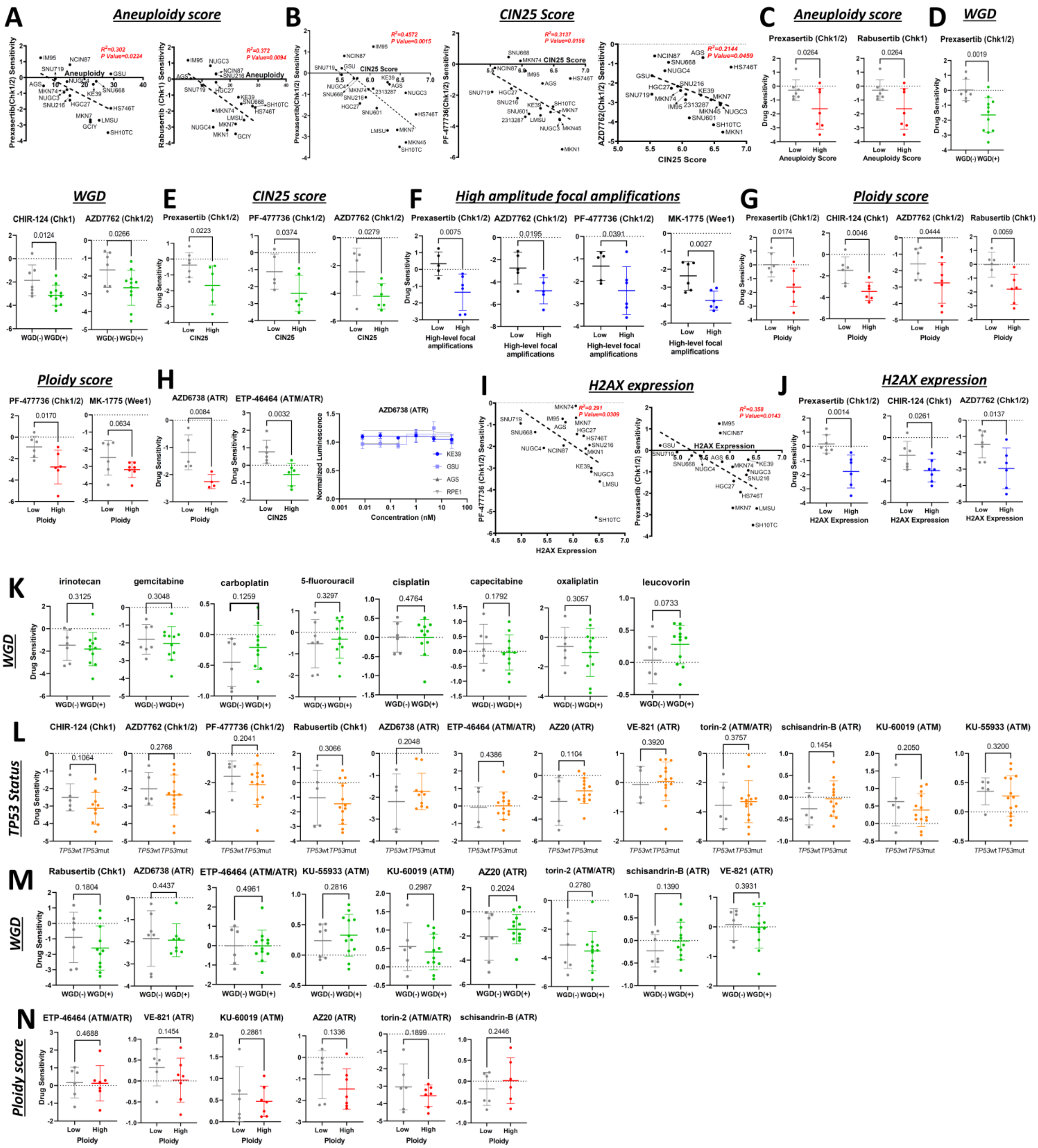

**Figure S4: Gastric cancer with high ploidy abnormalities and high *H2AX* expression are more sensitive to DDR pathway inhibitors (Chk1/2 and Wee1).**

A. Correlation plot of aneuploidy score with prexasertib (Chk1/2) and Rabusertib (Chk1/2) sensitivity (log2 fold change) for the CCLE gastric cancer cell lines; R squared value and p-value calculated by simple linear correlation analysis.

B. Correlation plot of CIN25 score with prexasertib (Chk1/2), PF-477736 (Chk1/2) and AZD7762 (Chk1/2) sensitivity for the CCLE gastric cancer cell lines; R squared value and p-value calculated by simple linear correlation analysis.

C. Chk1/2 inhibitors sensitivity between low aneuploidy score (grey) and high aneuploidy score (red) groups of gastric cancer cell lines available in BROAD institute PRISM repurposing drug screen dataset; Difference between the sensitivity (log2 fold change) is represented as the mean±S.D.; P-value calculated by unpaired t-test.

D. Chk1/2 inhibitors sensitivity between 'No WGD (0)' (grey) and 'with WGD (1/2)' (green) groups of gastric cancer cell lines available in BROAD institute PRISM repurposing drug screen dataset; Difference between the sensitivity (log2 fold change) is represented as the mean±S.D.; P-value calculated by unpaired t-test.

E. Chk1/2 inhibitors sensitivity between low CIN25 score (grey) and high CIN25 score (green) groups of gastric cancer cell lines available in BROAD institute PRISM repurposing drug screen dataset; the Difference between the sensitivity (log2 fold change) is represented as the mean±S.D.; P-value calculated by unpaired t-test.

F. Chk1/2 and Wee1 inhibitors sensitivity between 'low numbers of high amplitude focal amplifications' (grey) and 'high numbers of high amplitude focal amplifications' (blue) groups of gastric cancer cell lines available in BROAD institute PRISM repurposing drug screen dataset; Difference between the sensitivity (log2 fold change) is represented as the mean±S.D.; P-value calculated by unpaired t-test.

G. Chk1/2 and Wee1 inhibitors sensitivity between low ploidy score (grey) and high ploidy score (red) groups of gastric cancer cell lines available in BROAD institute PRISM repurposing drug screen dataset; Difference between the sensitivity (log2 fold change) is represented as the mean±S.D.; P-value calculated by unpaired t-test.

H. AZD6738 (ATR) sensitivity between low ploidy score (grey) and high ploidy score (red) groups of gastric cancer cell lines available in BROAD institute PRISM repurposing drug screen dataset; the Difference between the sensitivity (log2 fold change) is represented as the mean±S.D.; P-value calculated by unpaired t-test (left). ETP-46464(ATM/ATR) sensitivity between low CIN25 score (grey) and high CIN25 score (green) groups of gastric cancer cell lines available in the BROAD institute PRISM repurposing drug screen dataset; the Difference between the sensitivity (log2 fold change) is represented as the mean±S.D.; P-value calculated by unpaired t-test (middle). Dose-response curve of non-neoplastic cell line RPE1, Low replication-stress (grey)-AGS, and high replication-stress (blue)-KE39, GSU gastric cancer cell lines to the indicated concentrations of AZD6738 (ATR); Data presented as mean±S.D. of four culture replicates at each indicated dose (right).

I. Correlation plot of *H2AX* expression (log2(TPM+1)) with PF-477736 (Chk1/2) and prexasertib (Chk1/2) sensitivity (log2 fold change) for the CCLE gastric cancer cell lines; R squared value and p-value calculated by simple linear correlation analysis.

J. Chk1/2 inhibitors sensitivity between 'low *H2AX* expressing' (grey) and 'high *H2AX* expressing' (blue) groups (log2(TPM+1)) of gastric cancer cell lines available in BROAD institute PRISM repurposing drug screen dataset; Difference between the MK-1775 (Wee1) sensitivity (log2 fold change) is represented as the mean±S.D.; P-value calculated by unpaired t-test.

K. Chemotherapeutic agents sensitivity between 'No WGD (0)' (grey) and 'with WGD (1/2)' (green) groups of gastric cancer cell lines available in BROAD institute PRISM repurposing drug screen dataset; Difference between the sensitivity (log2 fold change) is represented as the mean±S.D.; P-value calculated by unpaired t-test.

L. DDR pathway inhibitors sensitivity between ‘*TP53*wt’ (grey) and ‘*TP53*mut’ (orange) groups of gastric Cancer cell lines BROAD institute PRISM repurposing drug screen dataset; Difference between the sensitivity (log2 fold change) is represented as the mean±S.D.; P-value calculated by unpaired t-test.

M. DDR pathway inhibitors sensitivity between 'No WGD (0)' (grey) and 'with WGD (1/2)' (green) groups of gastric cancer cell lines available in BROAD institute PRISM repurposing drug screen dataset; Difference between the sensitivity (log2 fold change) is represented as the mean±S.D.; P-value calculated by unpaired t-test.

N. DDR pathway inhibitors sensitivity between low ploidy score (grey) and high ploidy score (red) groups of gastric cancer cell lines available in BROAD institute PRISM repurposing drug screen dataset; the Difference between the sensitivity (log2 fold change) is represented as the mean±S.D.; P-value calculated by unpaired t-test.

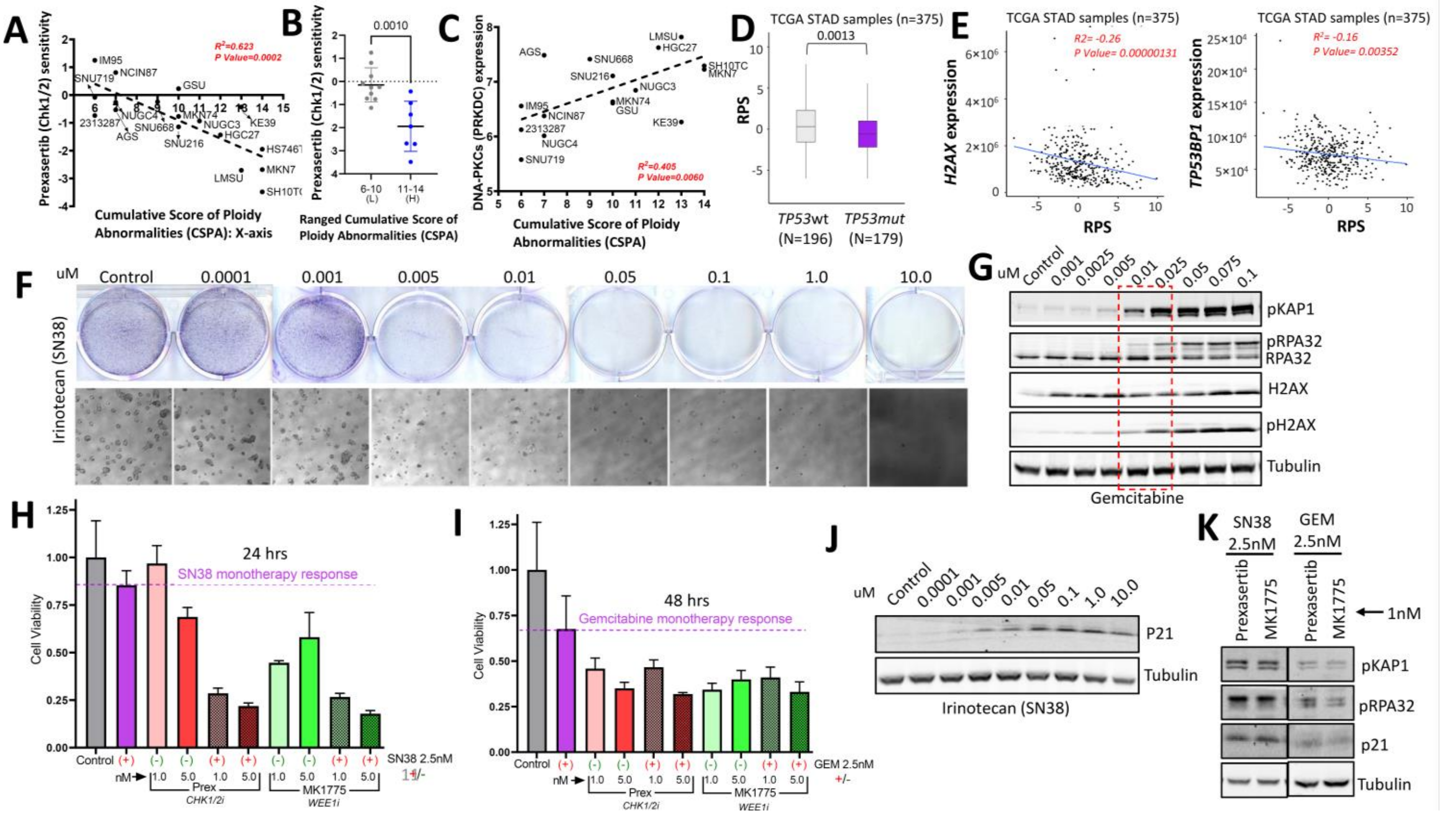


**Figure S5: Cumulative score of ploidy abnormalities predicts better prexasertib response, and irinotecan sensitizes DDR pathway inhibitors in gastric cancer cell lines; Defining the sensitivity and mechanism of DDR inhibitors/chemotherapeutic agents in gastric cancer cell lines with high replication stress.**

A. Correlation plot between 'Cumulative score of ploidy abnormalities' and 'prexasertib (Chk1/2)' sensitivity (log2 fold change) for the CCLE gastric cancer cell lines; R squared value and p-value calculated by simple linear correlation analysis.

B. prexasertib (Chk1/2) sensitivity between two sub-ranged 'Cumulative score of ploidy abnormalities' groups of gastric Cancer cell lines BROAD institute PRISM repurposing drug screen dataset; Difference between the prexasertib (Chk1/2) sensitivity (log2 fold change) is represented as the mean±S.D.; L=low, H=high; P-value calculated by unpaired t-test.

C. Correlation plot between 'Cumulative score of ploidy abnormalities' *and '*DNA-PKCs *(PRKDC)'* expression (log2(TPM+1)) for the CCLE gastric cancer cell lines; R squared value and p-value calculated by simple linear correlation analysis.

D. Recombination proficiency score (RPS) in '*TP53*wt' tumor, and '*TP53*mut' tumors from Stomach adenocarcinoma (STAD) cohort of TCGA.

E. Correlation plot between Recombination proficiency score (RPS), and *H2AX* /*TP53BP1* expression (FPKM) in patients from Stomach adenocarcinoma (STAD) cohort of TCGA.

F. Colony formation assay (top) showing irinotecan (SN38) dose-dependent sensitivity in KE39 cell line (high intrinsic replication stress) and representative bright-field images (below) for each concentration.

G. Immunoblot of replication stress/DDR markers and tubulin as the loading control in KE39 cell line (high intrinsic replication stress) treated with the indicated concentrations of gemcitabine (SN38).

H. Proliferation of the KE39 cell line (high intrinsic replication stress) treated with vehicle (grey), 2.5 nM of SN38 (magenta), prexasertib (red), prexasertib+SN38 (pattered red), MK1775 (green), MK1775+SN38 (pattered green); Light shade of color- 1.0nM of DDR inhibitor; Dark shade of color- 5.0nM of DDR inhibitor; All treatments for 24 hrs.

I. Proliferation of the KE39 cell line (high intrinsic replication stress) treated with vehicle (grey), 2.5 nM of gemcitabine (magenta), prexasertib (red), prexasertib+gemcitabine (pattered red), MK1775 (green), MK1775+gemcitabine (pattered green); Light shade of color- 1.0nM of DDR inhibitor; Dark shade of color- 5.0nM of DDR inhibitor; All treatments for 48 hrs.

J. Immunoblot of p21, and tubulin as the loading control in KE39 cell line (high intrinsic replication stress) treated with the indicated concentrations of irinotecan (SN38).

K. Immunoblot of pRPA32, pKAP1, p21, and tubulin as the loading control in KE39 cell line (high intrinsic replication stress) treated with combination therapy- SN38 (2.5nM)+prexasertib (1nM) and SN38 (2.5nm)+MK1775 (1nM), gemcitabine (2.5nM)+prexasertib (1nM), gemcitabine (2.5nm)+MK1775 (1nM).
